# Multi-modal representation of the size of space in the human brain

**DOI:** 10.1101/2023.07.24.550343

**Authors:** Jaeeun Lee, Soojin Park

## Abstract

To estimate the size of an indoor space, we must analyze the visual boundaries that limit the spatial extent and acoustic cues from reflected interior surfaces. We used fMRI to examine how the brain processes geometric size of indoor scenes when various types of sensory cues are presented individually or together. Specifically, we asked whether the size of space is represented in a modality-specific way or in an integrative way that combines multimodal cues. In a block-design study, images or sounds that depict small and large sized indoor spaces were presented. Visual stimuli were real-world pictures of empty spaces that were small or large. Auditory stimuli were sounds convolved with different reverberation. By using a multi-voxel pattern classifier, we asked whether the two sizes of space can be classified in visual, auditory, and visual-auditory combined conditions. We identified both sensory specific and multimodal representations of the size of space. To further investigate the nature of the multimodal region, we specifically examined whether it contained multimodal information in a coexistent or integrated form. We found that AG and the right IFG pars opercularis had modality-integrated representation, displaying sensitivity to the match in the spatial size information conveyed through image and sound. Background functional connectivity analysis further demonstrated that the connection between sensory specific regions and modality-integrated regions increase in the multimodal condition compared to single modality conditions. Our results suggest that the spatial size perception relies on both sensory specific and multimodal representations, as well as their interplay during multimodal perception.

## INTRODUCTION

Human perception relies greatly on vision (Pike et al., 2012) but it is rare that the human perception is achieved from one particular modality alone. We take in data from multiple sensory sources and make an integrated perception of the world (Ernst & Bülthoff, 2004; Stein & Meredith, 1993; van Atteveldt et al., 2014). The data can be complementary, where a piece of information can only be obtained from one sensory input, or they can be redundant, where the same information can be acquired from multiple sources (Noppeney, 2021). One example of redundant data is the representation of our surrounding environment—the salty sent of the ocean, the sound of waves crashing and bright blue colour of the water all indicate that we are at the beach. This wholesome representation is the result of our ability to incorporate sensory information from multiple sources. Then, how are the multiple sources of sensory information represented throughout various parts of the brain? Here we asked how the perception of spatial environment, specifically the geometric size of an indoor space, is represented in the human brain when visual and auditory cues are available. We also addressed how various parts of the brain represents the size of space from a single modal (visual or auditory) and multi-modal (visual and auditory) information.

Unbalanced amount of literature dedicated to topics of vision over other modalities such as auditory, tactile or olfactory (Gallace & Spence, 2014; Hutmacher, 2019) demonstrates the salience and importance of vision in understanding our perceptual world. Indeed, one can obtain significant amount of information from even a brief glance at a scene such as dominant depth and openness (Greene & Oliva, 2009) naturalness (Fei-Fei et al., 2007), 3D structure (Henderson et al., 2008), surface properties (Lowe et al., 2017) and even objects within it (Davenport & Potter, 2004). The rapid and sophisticated visual scene processing is supported by specialized neural substrates: the occipital place area (OPA; Dilks et al., 2013) encodes scene features that are important for visually guided navigation, such as path direction, first person motion, and distance to boundaries (Bonner & Epstein, 2021; Park & Park, 2020), the parahippocampal place area (PPA; Epstein & Kanwisher, 1998) encodes geometric spatial layout and semantic category information that are important for categorizing a place (Epstein, 2008), and the retrosplenial complex (RSC; Maguire, 2001) encodes mnemonic scene properties such as familiarity of scenes or map-based navigation information (Mitchell et al., 2018). In particular, evidence suggests that one of the geometric features encoded in these scene-specific regions is the spatial size of scenes (Park et al., 2015). For example, the PPA and RSC showed increased activity when scenes depicting large space (e.g., parking lots) were presented compared to scenes depicting small space (e.g., cockpit), and patterns of voxel activity in these regions were similar when two scenes shared similar size of space, even when semantic contents differed. These results imply that geometric size of space is one of the primary properties of visual scenes that are encoded in high-level scene-processing regions in the brain.

Auditory cues also play a role in representing the size of space. When soundwaves meet a hard surface like the wall of a room or an object, they are reflected, and we hear these reflections as echoes (Plack, 2018). Using this auditory feature, trained blind echolocation experts can discern the distance, size and motion of objects nearby (Stoffregen & Pittenger, 1995), and discriminate horizontal offsets of stimuli as small as 1.2° auditory angle (Teng et al., 2012) using self-generated sounds such as tongue clicks (Rojas et al., 2009). While previous literature shows that the sensory-motor union between vocalization and hearing must be well-established in order to accurately interpret echo information (Flanagin et al., 2017), those with unimpaired perceptual abilities can also gain geometric information of their surroundings through passive auditory resonant information. There is a very clear psychoacoustic perceptual difference between bouncing a ball in a small room and in an empty gym, implying that echo information is crucial for the estimation of the size of surrounding space. Supporting this idea, Teng and colleagues (Teng et al., 2017) used MEG to test the listening of various sounds revolved with different monaural room impulse response and showed that the reverberation response from surrounding space and the auditory source information are represented in different time and form within the brain (Teng et al., 2017). Specifically, the decoding of reverberant space peaked later (386 ms) than the source information (130 ms), while its neural response showed a more sustained pattern rather than transient.

Previous research has demonstrated how spatial size is represented by each visual (Park et al., 2015) and auditory (Teng et al., 2017) modality, but the multi-modal dynamic has not directly been addressed. One study integrated the neural dynamics of visual and auditory scene geometry processing by creating an MEG-fMRI spatio-temporal map (Cichy & Teng, 2017), but very few research examined how scene geometry is represented when two sensory stimuli are concurrently present. One study (Jung et al., 2018) used fMRI multi-voxel pattern classification for semantic scene categories (e.g., forest, city) when visual and auditory cues are independently and concurrently presented. They found both modality-specific coding and modality-independent coding of scene categories throughout different regions of the brain. While Jung et al., 2018 focused on semantic category representation of scenes, their results suggest a possibility that geometric size representation of scenes may also be represented by not only modality specific, but a multimodal, or even modality-independent ways.

In this study, we examine how spatial size of scenes is represented when visual, auditory, or both cues are presented. For those without severe sensory impairment, it is an effortless process to use both light information from the retina and acoustic wave energy reflected from a boundary surface reaching the ears to determine the size of their surroundings (Devore et al., 2009; Flanagin et al., 2017). Thus, the multimodal condition will allow us to examine how the brain represents the size of space when diverse modality cues are concurrently present, mimicking our natural daily perception.

We used fMRI multivoxel pattern analysis (MVPA) to test the neural representation of spatial size in indoor scenes from auditory and visual cues. Multiple regions-of-interests (ROIs) throughout the brain were examined to address how the spatial size representation varies between single-sensory and multisensory information processing. First, we predicted that sensory-specific cortical areas would represent spatial size of scenes based on each specific sensory cue. For example, when presented with Image-Only cue (Figure 1*A*), early to late visual processing regions such as V1, PPA, OPA and RSC would show accurate classification of spatial size from visual modality cue (Park et al., 2015); when presented with Sound-Only cue (Figure 1*A*), auditory processing regions such as Heschl’s gyri (transverse temporal gyri) would show accurate classification of spatial size from auditory modality cue (Di Salle et al., 2003).

**Figure 1.**
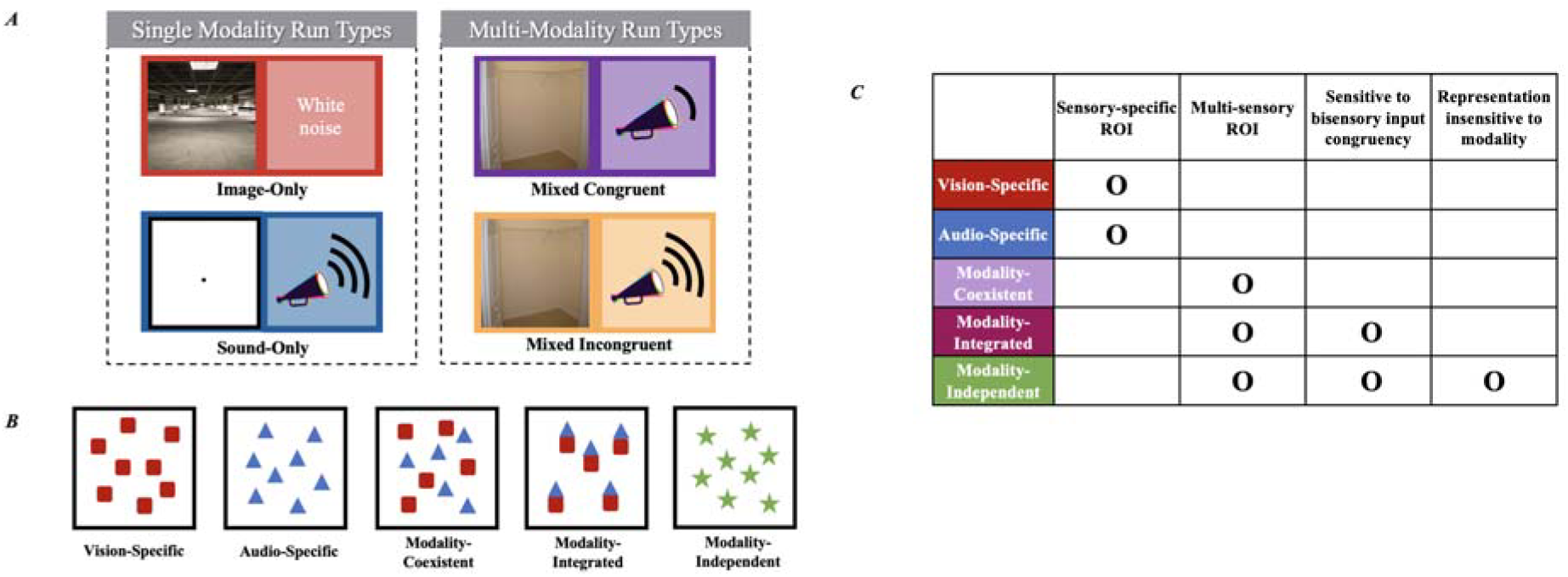
**A.** Illustration of the four experiment run types. During the single modality run types (Image-Only & Sound-Only), only either visual or auditory stimuli that indicated spatial size were presented. To keep the presentation environment similar across run types, a 2-second white noise audio clip was played per Image-Only trial, and a fixation dot was shown for 250ms per Sound-Only trial. Multi-modality run types includes Mixed Congruent and Mixed Incongruent, in which both visual and auditory stimuli were presented concurrently. **B.** Five possible models representing different types of spatial size information within the brain. Vision-Specific and Audio Specific ROIs would only contain information related to that modality. We introduce three types of Multi-sensory ROI models: Modality-Coexistent, Modality-Integrated, and Modality-Independent. **C.** Vision and Audio-Specific ROIs are categorized as sensory-specific. We can distinguish our three multi-sensory ROIs based on the following: A modality-coexistent region would contain both visual- and auditory-specific information, without generalization or additional sensitivity of the two information’s congruency. Modality-Integrated region also would contain information from the two modalities but would be sensitive to whether they are depicting the same spatial size. Modality-Independent, also known as supramodal model, would contain generalizable size representation that can cross-decode between visual and auditory information.

Next, we predicted that areas beyond the primary sensory regions may represent multimodal information. Several dynamics are possible in a voxel when processing more than one modality (Figure 1*B* & *C*). To test the nature of these representations, we concurrently presented images and sounds that either matched (Mixed-Congruent) or differed (Mixed-Incongruent) (Figure 1*A*) in terms of spatial size. We specifically tested three different possible models of how the brain may represent spatial size based on multimodal information: modality-coexistent, modality-integrated, and modality-independent. First, the modality coexistent model hypothesizes that both visual and auditory information to be present within a topographically adjacent region but not in an integrated form. An ROI with modality-coexistent representation would successfully decode spatial size from Image-Only, Sound-Only, and Mixed Congruent data but would be insensitive to the congruency of information coming from the two sensory sources. To test for sensitivity on bisensory input congruency (Figure 1*C*), we trained the decoder with Mixed-Congruent and tested on Mixed-Incongruent data. If the decoding accuracy is significant, this implies that the ROI is not sensitive to the congruency of spatial size information coming from image and sound and thus have modality-coexistent representation. On the other hand, if the decoding accuracy is not significant, this implies that the ROI is sensitive to the congruency of spatial size information conveyed through visual and auditory input and contains information that integrates size representations from both modalities. We call this latter case as the modality-integrated model. Lastly, a modality independent (supramodal) representation is when spatial size representation is exhibited as a form independent of the sensory source. The main difference between modality-independent and the other two representations is whether the generalization of the concept of spatial size exists. To test for the modality independent model, we trained the decoder with Image-Only data and tested with Sound-Only data, and vice-versa. If an ROI contains modality-independent representation of spatial size, then an ROI will show a significant cross-decoding between the two-modality data.

Including the Mixed-Incongruent condition, which is rare in harmonious natural settings, provides us an opportunity to unravel the different nature of multimodal representations. Using this condition, we also examined which modality’s information would prevail in processing spatial size when there is conflicting sensory input.

Complementing the hypothesis-driven ROI approach, we also employed whole-brain searchlight analysis to find modality-specific and multi-modal areas in the brain. Furthermore, we used background functional connectivity analysis to gain a better understanding about the dynamic nature of multisensory scene perception. While functional connectivity approach enables to uncover the dynamic between brain regions during rest period separated from task periods, the background connectivity approach enables to examine the spontaneous and intrinsic dynamic within the brain regions when participants are engaged by each condition, while controlling for the stimulus-evoked responses and noises removed (Al-Aidroos et al., 2012; Córdova et al., 2016; Norman-Haignere et al., 2012; Tompary et al., 2018). Specifically, we aimed to investigate the interplay between sensory regions when relevant sensory information is from a single source or multiple sources. We hypothesized that the connectivity between our multimodal and modality-specific ROIs would exhibit increased connectivity when both sound and visual stimuli were presented concurrently than when they were presented independently.

## MATERIALS AND METHODS

### Participants

Based on previous research (Dilks et al., 2013), we estimated our study to have a medium effect size (i.e., *d* = 0.7) with an alpha level of 0.05 and power level at 0.8. Our analysis revealed that a minimum of 15 people were needed to effectively test our study. Therefore, we recruited 17 right-handed subjects (11 females) aged between 19 and 29 (mean 25) years with normal or corrected-to-normal vision and with no hearing disabilities in the fMRI study for financial compensation. One subject’s data was excluded from the analysis due to extreme head movement. All subjects signed Informed Consent at the beginning of the experiment and the study protocol was approved by the Yonsei University Institutional Review Board.

### Stimuli

For visual stimuli, 14 color photographs of two scene sizes (large, small) were used. We collected images for both sizes from the stimuli set used in Park et al., 2015’s experiment 2. Small scenes included small closets and pantries, all of which were rated as space that would contain 1 to 2 people, and large scenes included empty parking garages and warehouses, all of which were rated as space that could hold around a thousand people. The low-level image statistics were controlled between the two sets so that the average amount of image spectral energy above the spatial frequency range of 10 cycles/image did not differ between the small and large scene sets (large: 26.4; small: 22.4; *t*_(6)_ = 0.76, *p =* 0.47). The visual stimuli were 500-by-500 pixels in resolution.

For auditory stimuli, three different sound clips (e.g., bouncing of a ball, tap of a pole, knock of wood) recorded in an anechoic chamber (from Teng et al., 2017) were used. To represent large indoor space, we used the reverberator system object in MATLAB 2020a Audio Toolbox to add reverberation to the original clips (decay factor of reverb tail: 0.3; density of reverb tail: 0.7; high-frequency damping: 0.004; default setting for other factors—wet-dry mix: 0.3; pre-delay for reverberation: 0; lowpass filter cutoff: 2000Hz). To represent small indoor space, the original clips with no reverberation were used. All sound clips were 2 seconds in length, sampling rate at 44.1 kHz, and loudness matched to -18 LUFS by ITU-R BS. 1770-3 algorithms in Adobe Audition. There was small difference in loudness across clips (bounce: 72.24 dB, tap: 71.80 dB, knock: 71.15 dB relative to the auditory threshold) but there was no difference in loudness between the spatial size categories.

To maintain participant’s attention, catch trials were randomly presented between stimuli once every block. For visual catch trials, a red frame (5-pixel width) appeared around the visual images; for auditory catch trials, a 440Hz pure tone was played randomly between sounds. The average percent correct across the four conditions was 99.36% (*Mdn* = 99.37%, range: 98.97 − 99.72%). Although the loudness and reverberation level are not directly based on exact volume of large scenes, the categorical distinction between small and large space sound clips were clear and all participants agreed in pre-experimental tests that the reverberation cue used in large-space sound match their perception of large empty spaces.

As described above, the visual and auditory stimuli representing small and large spaces were selected based on previous literatures, and pilot tests validated that participants were able to distinguish large and small spatial sizes from the stimuli set we used. To further ensure the appropriateness and discriminability of the stimuli, we additionally conducted an independent behavioral experiment using Amazon Mechanical Turk. A total of fifty participants were recruited and compensated for their involvement in the study. The first group of twenty-five participants were asked to judge whether the given sound originated from a small empty room suitable to fit approximately two people or a large empty room spacious enough to fit a thousand people. Another group of twenty-five participants were asked to judge the images the same manner. The average percentage of participants responding to the correct label was 91.42 % for images (*Mdn* = 92%, *SD* < 0.01 range: 84 − 100%) and 92% for sounds (*Mdn* = 90%, *SD* < 0.01 range: 88 − 100%). The results demonstrate a high level of consistency and agreement among participants in correctly categorizing the spatial sizes of the stimuli. The results of this post-hoc test provide further confirmation that the stimuli used in our study were reasonable representations of large and small spaces.

### MRI Experiment Procedure

There were four run types: Image-Only, Sound-Only, image & sound Mixed-Congruent, and image & sound Mixed-Incongruent. In the Image-Only and Sound-Only run types, participants either saw or heard the stimuli for the entire run. In the Mixed-Congruent and -Incongruent types, both visual and auditory stimuli were presented simultaneously, and the congruency was based on whether the spatial size of visual and auditory cues matched or not. The experiment consisted of 12 runs [each 4.53 min, 136 repetition time (TR)], 3 runs per run type, with 8 blocks per run. Each block represented scene size (large or small), and there were 4 blocks per scene size in a run. The order of blocks was randomized within a run. The order of runs was pseudo-randomized for each participant, with a constraint that the first six runs of the experiment were either Image-Only or Sound-Only run types, and the next six runs were the mixed run types. This pseudo-randomization assured participants to be familiar with the individual modality stimuli before the mixed run types. Due to technical difficulties, three participants only completed 8 out of 12 runs, resulting 2 runs per run type in these participants.

In the Image-Only run, a 16-second block consisted of seven scene images that represented either small or large space. The order of scenes within a block were randomized. Participants were instructed to press the button with their right forefinger if a red frame appeared around the stimulus image. These catch trials appeared once every block (14%). This somewhat orthogonal task was implemented based on previous literature on scene volume (Park et al., 2015) and object size (Konkle & Oliva, 2012). The inclusion of this task allowed the researchers to monitor participant’s attention and engagement throughout the run. Each image was shown for 250ms, then a blank screen was shown for 1750ms to complete 2 seconds. Inter-stimulus intervals (ISI) between the images ranged from 1770 to 1950ms so that the sum of their differences within a block did not exceed 2 seconds while keeping the overall length of a block to 16 seconds (Figure 2). To make sure that participants received both visual and auditory inputs during all run types, we played a two-second white noise clip while participants viewed visual scenes.

**Figure 2.**
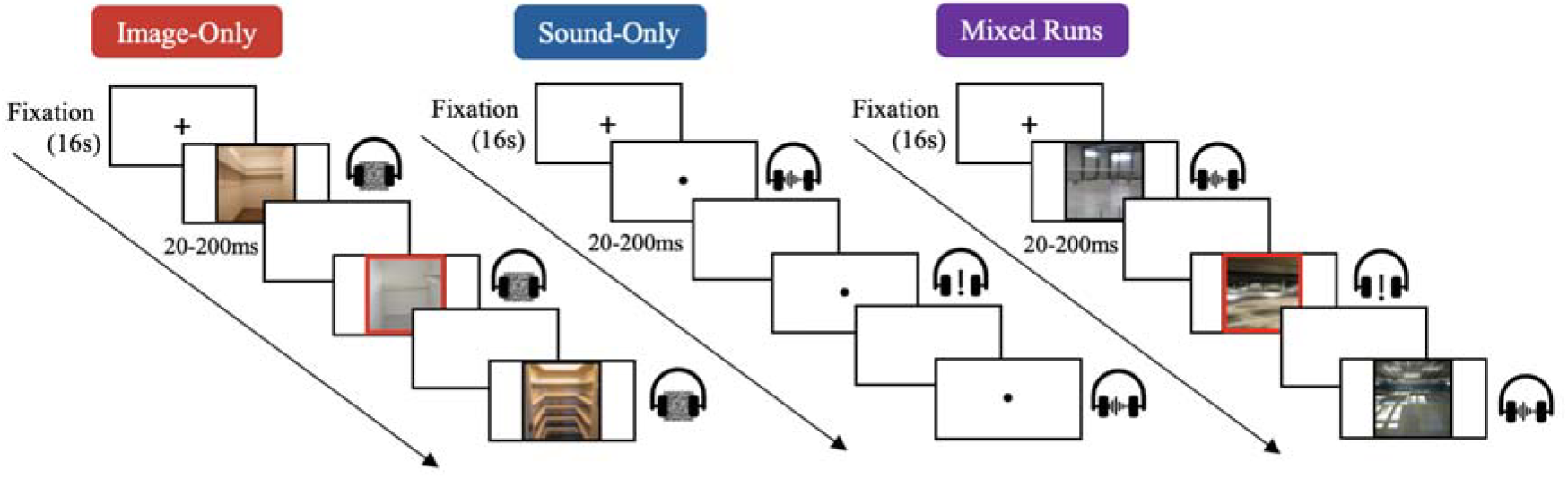
Example of stimuli presentation inside the MRI scanner. During Image-Only run types, participants were shown images of small or large-sized indoor scenes while white noise was played. Participants pressed a button when a red frame appeared around the stimulus image. For Sound-Only trials, a fixation dot was shown with the audio clips. Participants pressed a button when a pure tone stimulus played. During mixed run types, images and sounds were simultaneously presented and participants pressed a button when a red frame and a pure tone, which were always presented simultaneously, were presented.

In the Sound-Only run, each of the three sound clips were repeated twice in a randomized order and a two-second pure tone appeared as a catch trial (14%) within a block. Participants were instructed to press the button if a pure tone that is irrelevant to the stimulus was played. To make sure that participants received both visual and auditory inputs during all run types, a fixation cross appeared when the stimulus sound was played, and participants were asked to keep their eyes open even during the Sound-Only trials. All clips were played for 2 seconds with the ISI range between 20 and 200ms.

In the Mixed-Congruent run, images and sounds representing the same spatial size were presented simultaneously. In the Mixed-Incongruent run, two stimuli with different spatial sizes were presented. For example, if an image of a small closet was shown, a sound clip with large reverberation was played at the same time. The participants were instructed to think about the spatial size of the presented stimuli while pressing the button when both red frame and pure tone was presented. While rather artificial, these tasks allow the researcher to monitor whether the participants are paying attention to the visual and/or auditory stimuli. These tasks are commonly used in fMRI literature measuring perceptual processing of the stimulus set, separated from explicit high-level processing or mapping of the perceptual stimuli to motor responses. In the Mixed-Incongruent condition, they were asked to think about the spatial size that came to their minds first and were not restricted to pay attention to a specific modality. In both conditions, the catch trial appeared once every block (14%).

### MRI Data Acquisition and Preprocessing

MRI images were recorded using a 3T Phillips Ingenia CX Scanner located at Yonsei Medical Research Center. Functional images were acquired through a gradient echo, echo-planar sequence and high resolution, whole-brain structural images were also acquired before the functional scans for registration and anatomical localization. The experiment was presented via Matlab using the Psychtoolbox extensions (Brainard, 1997; Kleiner et al., 2007). Images were displayed invertedly on a monitor and were visible to the participants via a mirror mounted on the MR head coil (viewing distance: 1.5 m). Sounds were presented using the consoles and headphones from Nordic audio system and participants gave responses by pressing a button with their forefinger on the Lumina system button box. Functional data were analyzed using BrainVoyager QX software (Brain Innovation, Maastricht, Netherlands). Preprocessing included slice-timing correction, 3-D motion correction, temporal filtering with a high-pass filter at 3 cycles, and spatial smoothing with a Gaussian kernel with 4mm FWHM.

## DATA AND STATISTICAL ANALYSIS

### Regions of Interest

We incorporated published probabilistic maps (Wang et al., 2015) to localize the primary visual area (ventral V1) as well as parcels extracted using Group-Constrained Subject-Specific method (Julian et al., 2012) to localize high-level visual scene areas that encode spatial features such as geometry and mnemonic scene properties, including the OPA (Dilks et al., 2013), PPA (Epstein & Kanwisher, 1998), and RSC (Maguire, 2001). We selected these ROIs not only because many of them showed representation of geometric spatial size information in visual scenes, but also because we wanted to include visual ROIs that span across low to high level, to investigate the dynamic spatial size representation throughout the visual pathway. Because these ROIs utilized the Group-Constrained ROIs in a normalized space to start with, we defined these ROIs in each participants’ normalized MNI space. The anatomical ROIs in the temporal, parietal, and frontal lobes were extracted using the automated recon-all FreeSurfer reconstruction process (version 7.1.0; http://surfer.nmr.mgh.harvard.edu/) based on widely accepted cortical and subcortical atlas (Dale et al., 1999; Desikan et al., 2006; Fischl et al., 2004). These ROIs include: the primary auditory area (Heschl’s gyrus), middle temporal gyrus (MTG), superior temporal gyrus (STG), superior temporal sulcus (STS), angular gyrus (AG), superior parietal gyrus (SPG), intraparietal sulcus (IPS), medial frontal gyrus (MFG), superior frontal gyrus (SFG), and inferior frontal gyrus with pars opercularis (IFG opr), pars orbitalis (IFG orb), and pars triangularis (IFG tri). We chose the parietal and frontal ROIs based on previous works that investigated multi-modal and modality independent processing of visual and audio scene categories throughout the brain hierarchy. When scene images and sounds are decoded for its semantic category, the MTG, STG, STS, AG and SPG showed significant decoding of both visual and auditory based scene categories (Jung et al., 2018). Another study suggested that the IPS is involved in audiovisual processing (Calvert, 2001), so we chose to include this region as another parietal regions of interest. Given that the previous literature found modality independent coding of audio and visually presented scene categories in the pre-frontal cortex, we chose to include all anatomically defined frontal regions, from inferior, medial to superior frontal gyri. These anatomical ROIs were defined in the participant’s native space, and then normalized to MNI space. Although we defined each ROIs bilaterally, we combined the ROIs that exhibited no significant univariate differences between the hemispheres into a single ROI. Please refer to Figure 3 and Table 1 of the Appendix for details on the ROIs.

**Figure 3.**
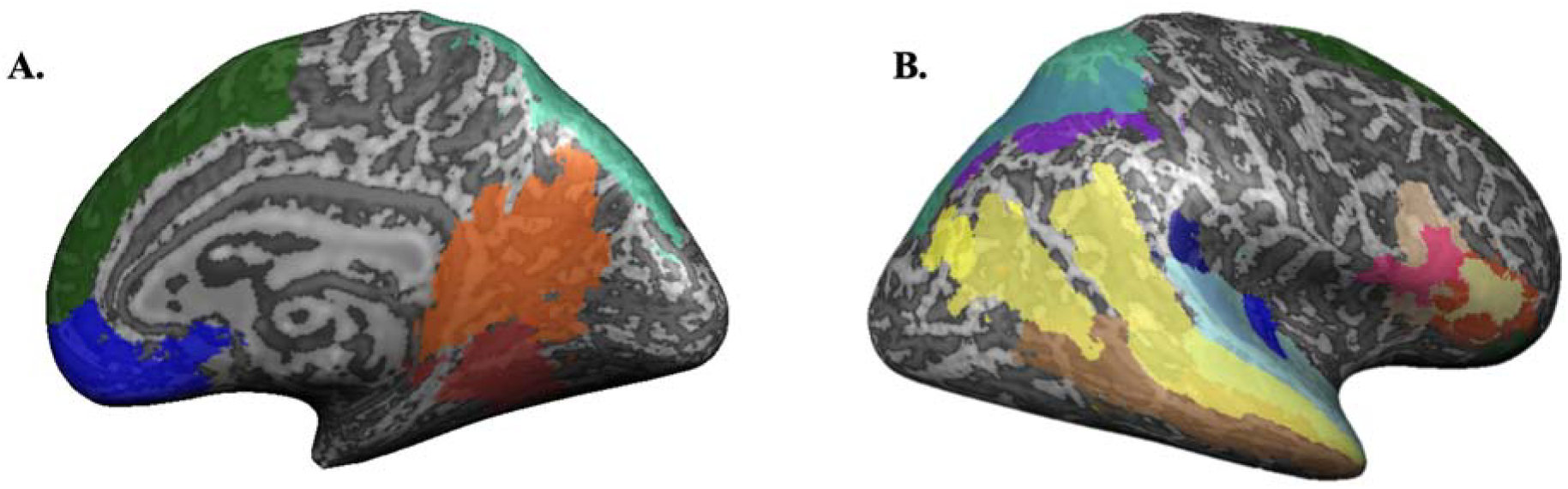
Anatomical and functional ROIs examined in this research. The areas are marked on the inflated surface of a participant’s brain. A. the medial surface view of the right hemisphere: SFG (dark green), MFG (dark blue), RSC (orange), PPA (red), SPG (cyan); B. the lateral surface of the right hemisphere: SPG (cyan), IPS (dark purple), STG (light blue), AG (dark blue—top), Heschl’s gyrus (dark blue—bottom), STS (yellow), MTG (brown), IFG opr (light beige), IFG tri (orange), IFG orb (faded yellow)

### ROI-based Multivariate Analysis

For multivariate pattern analysis (MVPA), no additional spatial or temporal smoothing was performed, and responses of each voxel were normalized into *z* scores to reduce the differences in signal across runs (Kamitani & Tong, 2005). These individual voxel activities of an ROI were averaged over time points within a block for each run condition.

We trained linear support vector machines using LIBSVM (Chang & Lin, 2011) in MATLAB (MathWorks, Inc.) to label the correct scene size label (small, large). Separate classifiers were used for each run type and ROI. For each classifier, leave-one-out cross validation method was used, in which all block data within the run except for one was used for training and the left-out run was tested. Multiple iterations of leave-one-run-out cross validation was performed so that all blocks contributed to both training and testing, and the averaged decoding accuracy from all iterations are reported. Specific hypotheses were tested using separate MVP classifiers (Figure 4). First, we performed decoding analysis in Image-Only runs and Sound-Only runs separately to test whether an ROI contained spatial size information specific to modality. Second, we trained and tested the decoder on Mixed-Congruent runs to determine which ROIs contained multisensory representations. Next, we performed MVPA using a decoder trained on Mixed-Congruent runs and tested on Modality-Incongruent runs to further dissociate modality-coexistent representation and modality-integrated representation. Given that Mixed-Incongruent condition has no correct answer in nature, we tested with different labels. For instance, if an image of a small closet was paired with a loud reverberation sound, the correct answer for image-based testing would be ‘small’, whereas for sound-based testing, it would be ‘large’. If an ROI contains modality-coexistent spatial size representation and insensitive to size congruency, it should be able to successfully decode either image- or sound-based testing, while modality-integrated ROIs would perform poorly in both cases. To supplement this analysis, we also directly compared decoding accuracies between Modality-Congruent and Modality-Incongruent runs within each ROI using *t*-tests, corrected for multiple comparisons using FDR (Benjamini & Hochberg, 1995). Lastly, we cross-decoded our single-modality runs, where the decoder was trained with Image-Only and test on Sound-Only data (and vice versa), to examine whether an ROI exhibited a modality-independent representation of spatial size.

**Figure 4.**
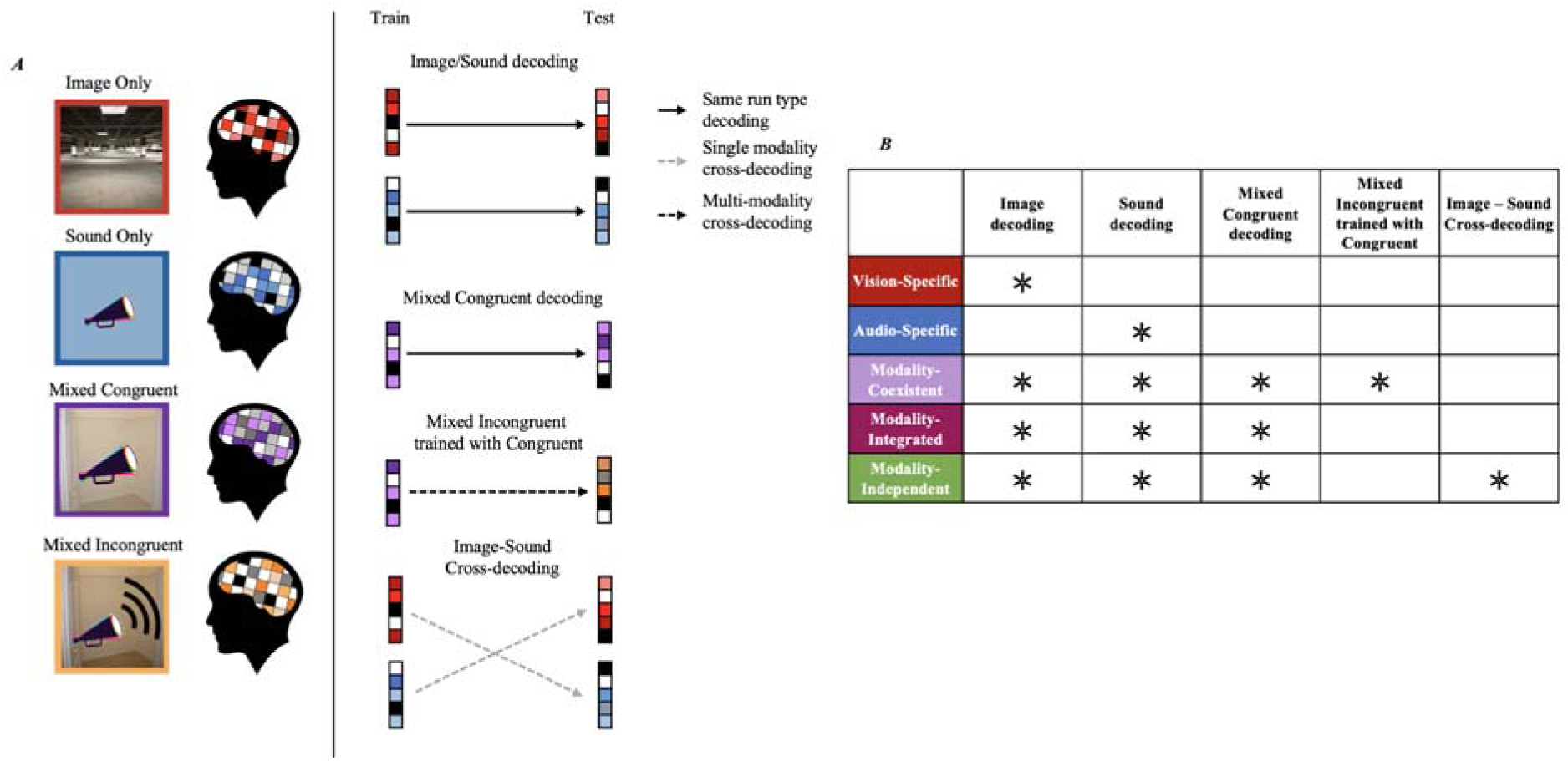
**A.** Illustration of the MVPA decoding process. Separate decoders were trained for each run type and ROI. Each run type is color coded to represent Image-Only (red), Sound-Only (blue), Mixed Congruent (purple), Mixed Incongruent (orange) conditions. The straight black arrows indicate classifiers in which the training and testing of the decoder is from the same run type (i.e., training the classifier with Image-Only data and testing it on data with the same condition and this also applies to Sound-Only and Mixed Congruent). The dotted black line represents the case where the classifier was trained with Mixed Congruent condition and tested with Mixed Incongruent condition. The dotted grey line indicates classifiers in which the training data and the testing data are from different run types (cross-decoding, i.e., training the classifier with Image-Only data and testing on Sound-Only data). **B.** Table summarizing different ROI types and decoding process. The asterisks each row show the decoding conditions that must be satisfied.

In all of the statistical testing of our decoding results, we used a nonparametric permutation approach (Meyyappan et al., 2021; Stelzer et al., 2013). The null distribution was generated using a permutation test by shuffling the run labels (small vs. large spatial size) 100 times for each participant. This process yielded 100 chance accuracies per participant. Next, we used a bootstrapping method to select one accuracy value from the pool of chance accuracies 100,000 times (Elli et al., 2019; Stelzer et al., 2013). By using this approach, we generated an empirical null distribution for each ROI and condition across participants. The mean of this distribution was then used to obtain the *t*-statistics. In addition, to account for multiple comparisons, the significance of the *t*-test results was corrected using false discovery rate (FDR) technique (Benjamini & Hochberg, 1995).

### Whole-brain Searchlight Analysis

To investigate any cortical regions with a modality-independent representation that was outside our predefined ROIs, we performed a roving searchlight analysis (Kriegeskorte et al., 2006) using a 3mm radius spherical neighborhood around each voxel in the cortical sheet. The pattern of the voxels in that neighborhood went through the same leave-one-out cross-validation performed using a linear SVM classifier, same as the ROI-based multivariate analysis. Group analysis was performed by overlaying each participant’s decoding accuracy maps to MNI space, the atlas to which all the subjects’ brains were coregistered. To identify the significant voxels at the group level, we performed one-tailed *t* tests on whether the sphere around each searchlight center had decoding accuracy above chance level.

### Background Functional Connectivity Analysis

We assessed the background connectivity between modality specific and multimodal areas to gain some insight into how dynamics of multimodal ROIs change as the modality of input changes. To do so, we examined correlations between our predetermined regions of interests independent of the evoked responses to individual stimuli. This analysis reflects the BOLD correlations between two seed regions while maintaining a certain task, but with the mean evoked response removed (Al-Aidroos et al., 2012; Córdova et al., 2016; Norman-Haignere et al., 2012; Tompary et al., 2018). The preprocessed data of each participant was fit using a finite impulse response (FIR) model because it is free from assumptions regarding the shape of the hemodynamic response. Our general linear model consisted of 24 constant height candlestick regressors that represented each time point within a single block after the onset of the stimulus. As a result, any response that took place after the onset would be accounted for by at least one of the 24 predictors, allowing us to capture the temporal dynamic of the neural responses in a more precise manner (Norman-Haignere et al., 2012). We extracted the residuals from each ROI from this model and these residuals were then scrubbed of nuisance variance using another GLM model. This new model includes a global mean of the BOLD time course over all the voxels, the means of four white matter, four ventricle seeds and six motion correction parameters obtained from preprocessing (Figure 5). We searched for spheres of 3 mm radius, each from the left and right anterior and posterior regions of the centrum semiovale for white matters and ventricle in order to determine the eight seeds (Hayasaka, 2013). Out of all the spheres found, we chose the most anterior or posterior ones for each participant. However, a 2mm radius sphere was fit for one participant’s ventricle seeds because we could not find a 3mm-radius sphere.

**Figure 5.**
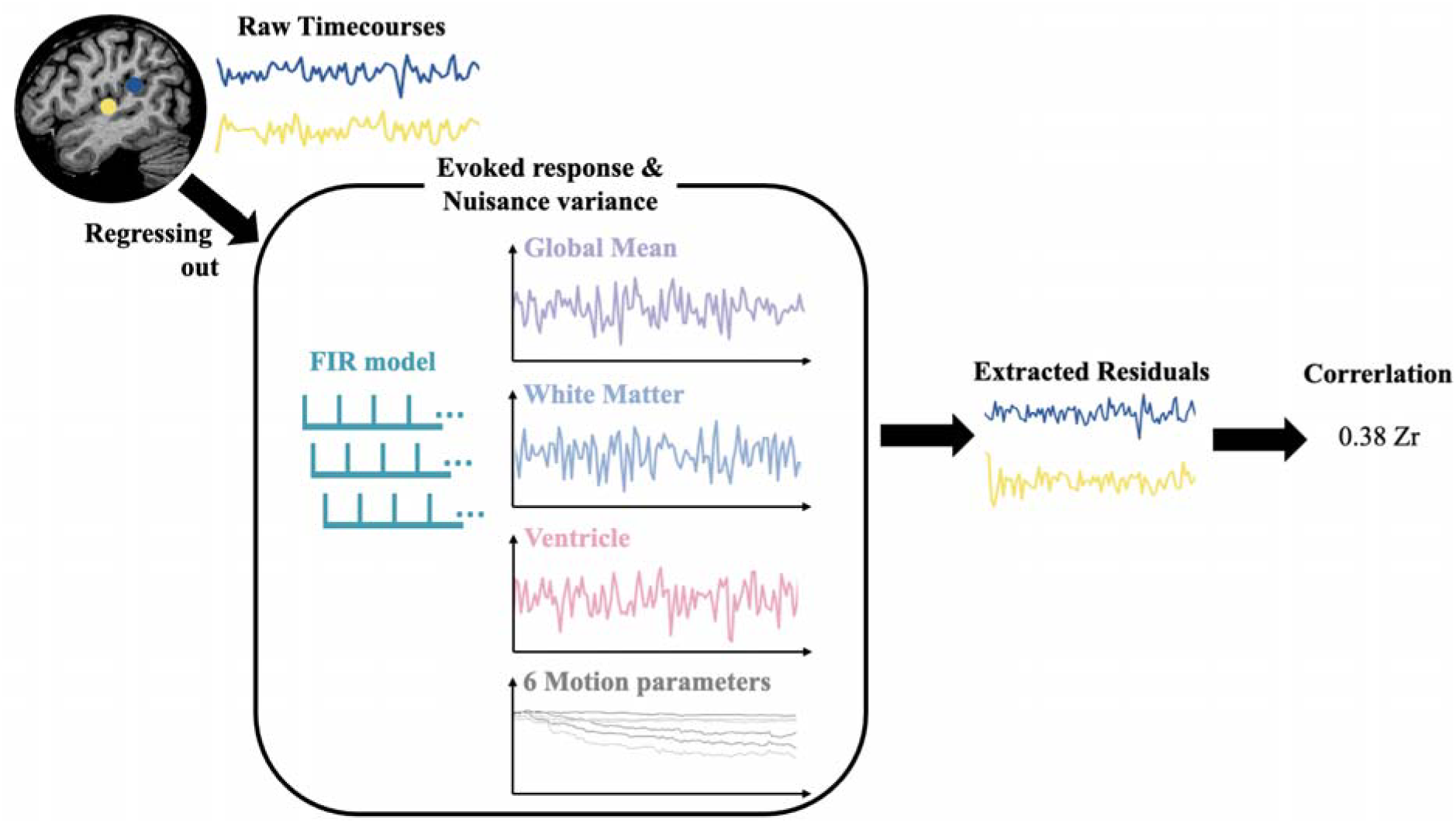
A schematic of background functional connectivity analysis. After first extracting the raw time courses of our selected ROIs (marked as blue and yellow circles on the brain), we regressed out the mean evoked response of conditions using the FIR model, then removed nuisance variables such as the six motion parameters, mean of the four white matter seeds, mean of the four ventricle seeds, and the global mean of the whole brain signals. With these extracted residual time courses, we calculated the correlation to obtain a measure of the background connectivity index.

The residual data from the Mixed-Incongruent run type was utilized to determine the peak voxel in each ROI. Since the aim of the functional connectivity analysis was targeted to examine whether there is an increase in correlation for modality integrated stimuli (Mixed-Congruent) compared to single modality stimuli (Image-Only, Sound-Only), we utilized the left out condition to eliminate any bias in selecting seeds relevant to the run conditions being tested. Given our objective of observing the correlation dynamic between single- and multi-modal ROIs, we incorporated five modality-specific ROIs (V1, right OPA, PPA, RSC and Heschl’s gyri) and two multi-modal regions (AG, and the right IFG opr). To ensure focused analysis, spheres with a radius of 4mm were created around the peak voxels within these ROIs. These spheres served as seeds for the correlation analysis, instead of the whole ROI. Defining seed ROIs based on spheres around peak activity has advantages of increased specificity to enable a more targeted examination and reduction of noise which is particularly important in the background functional connectivity analysis. From these seeds, we extracted the residual data for the rest of the run conditions, Image-Only, Sound-Only, and Mixed-Congruent. Afterwards, the residual mean time courses were correlated with each other using Pearson correlation analysis. The correlation coefficients were then transformed into *z* scores using Fisher’s *r*-to-*z* transformation.

## RESULTS

### ROI-based Analysis

#### Decoding single-modality data

First, to examine which brain areas encode the spatial size information of a specific modality, we performed MVPA for each of our ROIs on run types consisting only of single sensory stimuli (Image-Only and Sound-Only runs). Modality specificity of an ROI would indicate that size of an indoor space can be sufficiently represented from a single source of sensory input. For size decoding using Image-Only runs, we found significant scene size classification in the early visual areas in the ventral visual pathway (V1: mean: 58.3%, *t*_(15)_ = 2.89, *q* = 0.01, *d* = 0.72), and most of the scene-specific areas except for the left PPA. The areas include the right PPA (mean: 60.3%, *t*_(15)_ = 3.51, *q* < 0.01, *d* = 0.88), right and left OPA (right OPA: mean: 59.6%, *t*_(15)_ = 3.14, *q* = 0.01, *d* = 0.79; left OPA: mean: 60.5%, *t*_(15)_ = 3.51, *q* < 0.01, *d* = 0.87), and right and left RSC (right RSC: mean: 54.3%, *t*_(15)_ = 1.91, *q* < 0.05, *d* = 0.48; left RSC: mean: 56.9%, *t*_(15)_ = 2.55, *q* < 0.05, *d* = 0.64). Interestingly, the primary auditory area in the Heschl’s gyrus could also accurately classify spatial size with visual cues (mean: 58.6%, *t*_(15)_ = 2.37, *q* < 0.05, *d* = 0.448). For size decoding using Sound-Only runs, the primary auditory area could successfully represent spatial size using auditory cues (Heschl’s gyrus: mean 66.1%, *t*_(15)_ = 5.15, *q* < 0.01, *d* = 1.282). Interestingly, some of the visually specific brain areas also accurately classified spatial size with auditory cues as well (Appendix Table 2). For example, spatial size in Sound-Only runs were significantly classified in V1 (mean: 56.9%, *t*_(15)_ = 2.97, *q* = 0.01, *d* = 0.74) and all of the scene-specific areas including bilateral RSC (right RSC mean: 61.1%, *t*_(15)_ = 3.79, *q* < 0.01, *d* = 0.71; left RSC 61.3%, *t*_(15)_ = 3.32, *q* < 0.01, *d* = 0.82).

The asymmetrical representation between visual and auditory cues suggest that visually presented scene size information is represented more specifically in the high-level category specific regions within the ventral visual stream (PPA and OPA) than the primary visual area, while auditory scene-size information is represented specifically as early as in primary auditory cortex. Beyond sensory-specific regions, our temporal and parietal areas all showed significant above-chance accuracy for both visual run types and auditory run types. These include the MTG (Image-Only mean: 61.6%, *t*_(15)_ = 4.12, *q* < 0.01, *d* = 1.03; Sound-Only: 61.7%, *t*_(15)_ = 3.17, *q* = 0.01, *d* = 0.79), STG (Image-Only mean: 61.7%, *t*_(15)_ = 3.64, *q* < 0.01, *d* = 0.91; Sound-Only: 65.2%, *t*_(15)_ = 4.61, *q* < 0.01, *d* = 1.15), STS (Image-Only mean: 60.4%, *t*_(15)_ = 4.14, *q* < 0.01, *d* = 1.04; Sound-Only: 59.8%, *t*_(15)_ = 2.95, *q* = 0.01, *d* = 0.74), AG (Image-Only mean: 58.3%, *t*_(15)_ = 3.57, *q* < 0.01, *d* = 0.89; Sound-Only: 65.9%, *t*_(15)_ = 6.42, *q* < 0.01, *d* = 1.61), SPG (Image-Only mean: 61.5%, *t*_(15)_ = 3.66, *q* < 0.01, *d* = 0.92; Sound-Only: 59.9%, *t*_(15)_ = 4.27, *q* < 0.01, *d* = 1.06), and IPS (Image-Only mean: 60.7%, *t*_(15)_ = 3.59, *q* < 0.01, *d* = 0.90; Sound-Only: 58.7%, *t*_(15)_ = 2.84, *q* = 0.01, *d* = 0.71). Most of the ROIs from the frontal cortex also showed above-chance decoding accuracy for both Image-Only and Sound-Only run conditions including the bilateral IFG opr (right Image-Only mean: 58.6%, *t*_(15)_ = 3.02, *q* = 0.01, *d* = 0.76; Sound-Only; 56.8%, *t*_(15)_ = 2.73, *q* = 0.01, *d* = 0.68; left Image-Only mean: 60.4%, *t*_(15)_ = 3.93, *q* < 0.01, *d* = 0.98; Sound-Only mean: 57.7%, *t*_(15)_ = 3.93, *q* = 0.01, *d* = 0.81), IFG orb (right Image-Only mean: 55.9%, *t*_(15)_ = 2.49, *q* < 0.05, *d* = 0.62; Sound-Only; 57.9%, *t*_(15)_ = 2.57, *q* = 0.01, *d* = 0.64; left Image-Only mean: 56.4 %, *t*_(15)_ = 3.30, *q* = 0.01, *d* = 0.83; Sound-Only mean: 61.1%, *t*_(15)_ = 3.14, *q* = 0.01, *d* = 0.78), IFG tri (right Image-Only mean: 57.6%, *t*_(15)_ = 2.84, *q* = 0.01, *d* = 0.71; Sound-Only; 60.4%, *t*_(15)_ = 3.41, *q* < 0.01, *d* = 0.85; left Image-Only mean: 61.8%, *t*_(15)_ = 3.40, *q* = 0.01, *d* = 0.85; Sound-Only mean: 57.8%, *t*_(15)_ = 2.39, *q* < 0.05, *d* = 0.60), right SFG (Image-Only mean: 62.6%, *t*_(15)_ = 4.33, *q* < 0.01, *d* = 1.08; Sound-Only: 62.8%, *t*_(15)_ = 3.45, *q* < 0.01, *d* = 0.86) and left SFG (Image-Only mean: 65.6%, *t*_(15)_ = 4.81, *q* < 0.01, *d* = 1.20; Sound-Only: 61.5%, *t*_(15)_ = 3.18, *q* = 0.01, *d* = 0.80). These regions show significant decoding for both modalities, suggesting that multi sensory representation of scene space may exist in these areas of the temporal and parietal cortices. We test such possibilities more explicitly by decoding multi-modality data next.

#### Decoding multi-modality data

For the Mixed-Congruent run type, we concurrently presented both scene images and sounds that matched in spatial size. We asked if the regions that could decode information coming from the two single modalities could also decode multi-modal information (Mixed-Congruent run type). Such regions would indicate a multimodal representation of spatial size, where neurons responsible for image and sound cues coexist in the voxels.

Training and testing the support vector machine with Mixed-Congruent data revealed significant classification accuracy in the primary visual area (V1: mean 57.7%, *t*_(15)_ = 3.88, *q* < 0.01, *d* = 0.97), Heschl’s gyrus (mean 71.1%, *t*_(15)_ = 5.06, *q* < 0.05, *d* = 1.26), temporal, and parietal cortex. Specifically, in temporal and parietal cortices, significant regions of interests include STG (mean: 64.2%, *t*_(15)_ = 5.59, *q* < 0.01, *d* = 1.40), AG (mean: 62.5%, *t*_(15)_ = 3.92, *q* < 0.01, *d* = 0.98), and IPS (mean: 59.1%, *t*_(15)_ = 3.40, *q* = 0.01, *d* = 0.85). In the prefrontal regions, all subregions of the right IFG were significant (pars opercularis mean: 56.9%, *t*_(15)_ = 4.38, *q* < 0.01, *d* = 1.10; pars orbitalis mean: 56.4%, *t*_(15)_ = 2.57, *q* < 0.05, *d* = 0.64; pars triangularis: 58.5%, *t*_(15)_ = 3.95, *q* < 0.01, *d* = 0.99), as well as bilateral SFG (right mean: 59.6%, *t*_(15)_ = 3.95, *q* < 0.01, *d* = 0.98; left: 60.4%, *t*_(15)_ = 3.23, *q* = 0.01, *d* = 0.81). Considering that these ROIs also showed significant decoding of spatial sizes when using single modality data, the significant decoding of Mixed-Congruent runs suggest that these ROIs contain multimodal representation of scene size.

Do these multi-modal ROIs simply contain multi-modal information in a topographically clustered way (modality-coexistent), or contain integrated information between the two modalities (modality-integrated)? To specifically separate the two possibilities, we tested above multi-modal ROIs for sensitivity to multi modal congruency by training the decoder with Mixed-Congruent data and testing on Mixed-Incongruent data. Please note that, given the nature of Mixed-Incongruent run, the analysis is divided into two: one tested with Mixed-Incongruent data labeled based on image, and another tested with Mixed-Incongruent data labeled based on sound. If the decoding results are significant when trained with Mixed-Congruent data and tested with Mixed-Incongruent data, in either one of the labels, it suggests that the ROI is insensitive to the congruency between the two modality and suggests that the ROI is simply modality coexistent in nature. By further analyzing whether the decoding results are significant for one label-based analysis but not the other, we can further infer which sensory modality prevails when visual and auditory inputs co-exist in an ROI but convey contradicting information. Specifically, if the decoding accuracy of an ROI is significant when the testing label was based on image but not when based on sound, we can infer that the nature of multimodal ROI is biased by preference to visual modality over auditory when the two are disharmonious. On the other hand, if the decoding results are not significant for any types of labels, it suggests that the ROI is sensitive to the congruency between visual and auditory inputs and contains modality-integrated representation.

When we tested the decoder with Mixed-Incongruent runs labeled based on images, most of the multi modal ROIs showed significant decoding accuracy. Examples include V1 (mean: 63.3%, *t*_(15)_ = 6.73, *q* < 0.01, *d* = 1.69), STG (mean: 52.7%, *t*_(15)_ = 2.55, *q* < 0.01, *d* = 0.75), IPS (mean: 61.5%, *t*_(15)_ = 4.75, *q* < 0.01, *d* = 1.19), right IFG orb (mean: 57.2%, *t*_(15)_ = 3.02, *q* < 0.05, *d* = 0.64) and bilateral SFG (right mean: 55.7%, *t*_(15)_ = 4.06, *q* < 0.01, *d* = 1.01; left mean: 55.7%, *t*_(15)_ = 2.93, *q* = 0.01, *d* = 0.73). These ROIs satisfy our definition of modality-coexistent (Figure 4 B). However, this was not the case when the labels were based on sounds. None of our preselected ROIs showed significant decoding accuracies for this condition. This suggests that while these ROIs contain modality-coexistent representation, the ROIs have preference for spatial size information conveyed through visual input compared to auditory, when there is conflict of information between the two.

Importantly, AG and the right IFG opr did not show any significant results when tested with either the image-based or sound-based Mixed-Incongruent runs. Critically, these are the regions that previously showed significant decoding when trained and tested with Mixed-Congruent runs. This suggests that, while these ROIs contain multi-sensory information, the representation do not transfer to Mixed Incongruent runs, suggesting that these regions are sensitive to the congruency between the two modalities. In other words, the multi-modal representation in these ROIs is modality-integrated in nature, containing information about the congruency between the two modalities, beyond mere co-existence of the two.

#### Cross-decoding between Image- and Sound-Only run types

To ask whether there are any regions that represent spatial size information independent of information restricted from a certain sensory input, we tested cross-decoding between the Image-Only and Sound-Only run types. However, we could not find any region that could successfully decode any cross-analysis cases. In other words, no modality-independent representations were found in our ROIs. Therefore, we explored beyond our ROIs to search for modality-independent representation by using whole brain searchlight analysis.

### Whole-brain Searchlight Analysis

While the ROI-based analyses revealed that functionally and anatomically pre-determined regions of interests did not display modality-independent representation, there may be regions outside of predetermined ROIs that could have representations of indoor spatial size. We used searchlight analysis to find such areas that may show modality-independent, or cross-modal representations. Specifically, we performed four different classifiers in the whole-brain searchlight analysis: Image-Only decoding, Sound-Only decoding, cross-decoding trained with Image-Only and tested with Sound-Only, and cross-decoding trained with Sound-Only and tested with Image-Only. A region with modality-independent representation should show above-chance level decoding accuracy for all four analyses above.

First, we found voxels overlapping with the right OPA and PPA to represent spatial size information from Image-Only runs (Figure 6*B*), consistent with our ROI-based classification results (*p < 0.001, cluster threshold = 30)*. Next, we found voxels in the STG, and the Heschl’s gyrus in both left and right hemispheres represented spatial size information from Sound-Only runs (Figure 6*C*). Finally, confirming our ROI-based multivoxel pattern analysis, the searchlight analysis did not reveal any significant cross decodable voxels between Image-Only and Sound-Only conditions. When we trained the decoder with Image-Only and tested with Sound-Only data, no voxel cluster made it above the threshold, and neither was the case of training with Sound-Only runs and testing with Image-Only runs. These results confirm that the size of space using both visual and auditory cues is represented in a multimodal way in the brain, rather than modality-independent (supramodal) way.

**Figure 6.**
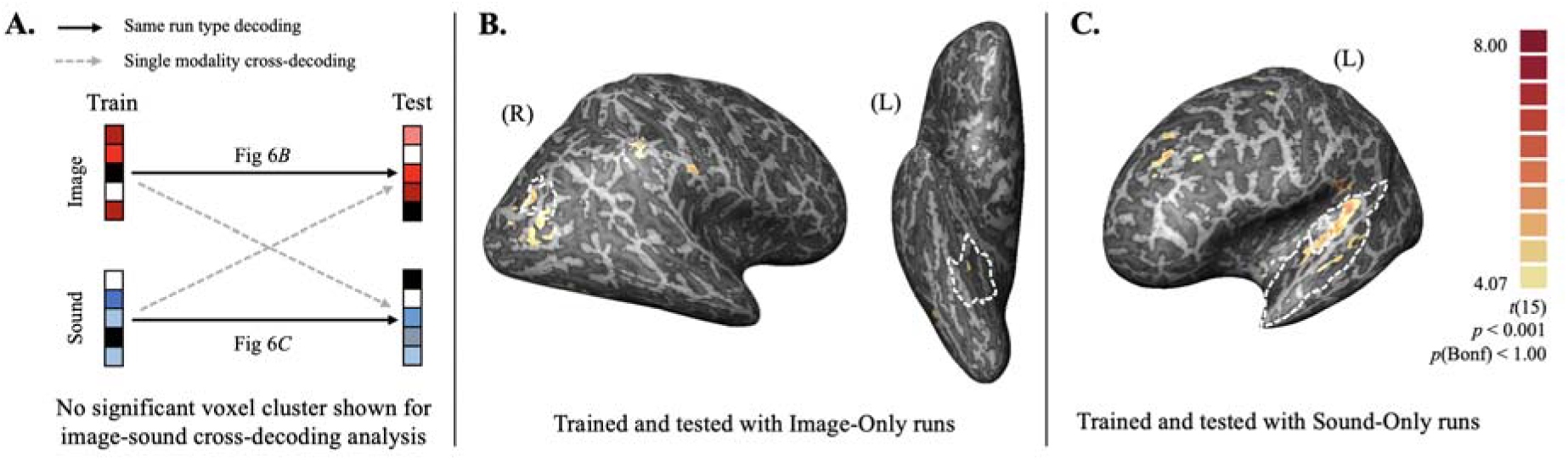
Results of whole-brain searchlight analysis overlapped with our predefined ROI. **A.** Diagram of the analysis. Cross-decoding between Image-Only and Sound-Only runs (marked with dotted gray line) revealed no significant voxel cluster that made above our threshold. **B.** Regions with significant decoding accuracy for Image-Only runs. Here, the decoder was trained with Image-Only data and tested on the Image-Only data not included in training. White dotted lines on the left represent the right OPA and dotted lines on the left represent the right PPA. **C**. Regions in the left hemisphere with significant decoding accuracy for Sound-Only runs. Likewise, the decoder was trained with Sound-Only data and tested on the Sound-Only data not included in training. Outer white dotted lines depict the STG and inner lines depict the Heschl’s gyrus.

### Background Functional Connectivity Analysis

We used background functional connectivity analysis to investigate the connection dynamic between low-level sensory regions and multimodal regions in response to audio, visual, or audio-visual stimuli. One model of the dynamic suggests that the multisensory processing found in multimodal regions is based on the summation of feedforward information from multiple sensory regions. An alternative model is that there are more complicated dynamics in multimodal brain regions involving additional interaction among ROIs. In this case, we anticipate finding stronger connectivity between modality-specific and modality-integrated ROIs in Mixed Congruent runs compared to single sensory runs. We hypothesized that the different run types would engender distinct states of background connectivity between our ROIs. Moreover, based on the interesting ROI-based MVPA finding of V1 demonstrating significant decoding accuracy in the Sound-Only run type, we also examined whether this observation was reflected in the connectivity between V1 and multimodal areas. Specifically, we analyzed the background connectivity between five sensory-specific ROIs (V1, right OPA, PPA, RSC and Heschl’s gyri) and two modality integrated regions (AG, and the right IFG opr). Since our goal was to examine whether there is an increase in correlation for multimodal stimuli compared to single modality stimuli, we focused on analyzing the connectivity of modality-integrated ROIs than modality-coexistent ROIs.

To find the background connectivity in each region, we extracted the residual activation of our predetermined ROIs by leaving out the nuisance and mean evoked responses from our neural data. These regions’ residual time courses were correlated, and this correlation coefficient was converted to a *z* score using Fisher’s *r*-to-*z* transform, resulting three 77 correlation matrices for each run type (Image-Only, Sound-Only, and Mixed-Congruent). To examine the correlation dynamic closer, we first contrasted the Mixed-Congruent with Image-Only and Sound-Only data. This analysis reflects whether background connectivity between the two ROIs that are being compared gets stronger when multisensory cues rather than single cues, are presented. Among our ROIs, we found relatively higher correlation between the single modality areas (right OPA, PPA and Heschl’s gyrus) and multimodal areas when bisensory cues were shown, so we further ran one-tailed *t* tests corrected using Benjamini-Hochberg method (Benjamini & Hochberg, 1995) to investigate its statistical significance. Results showed that the background connectivity between OPA and AG was significantly stronger (*t*_(15)_ = 2.41, *q* < 0.05, FDR corrected) for Mixed-Congruent than Image-Only run types. The Heschl’s gyrus showed marginally significant difference between Mixed-Congruent and Sound Only run types with AG (*t*_(15)_ = 1.90, *q* =0.08, FDR corrected) (Figure 7).

**Figure 7.**
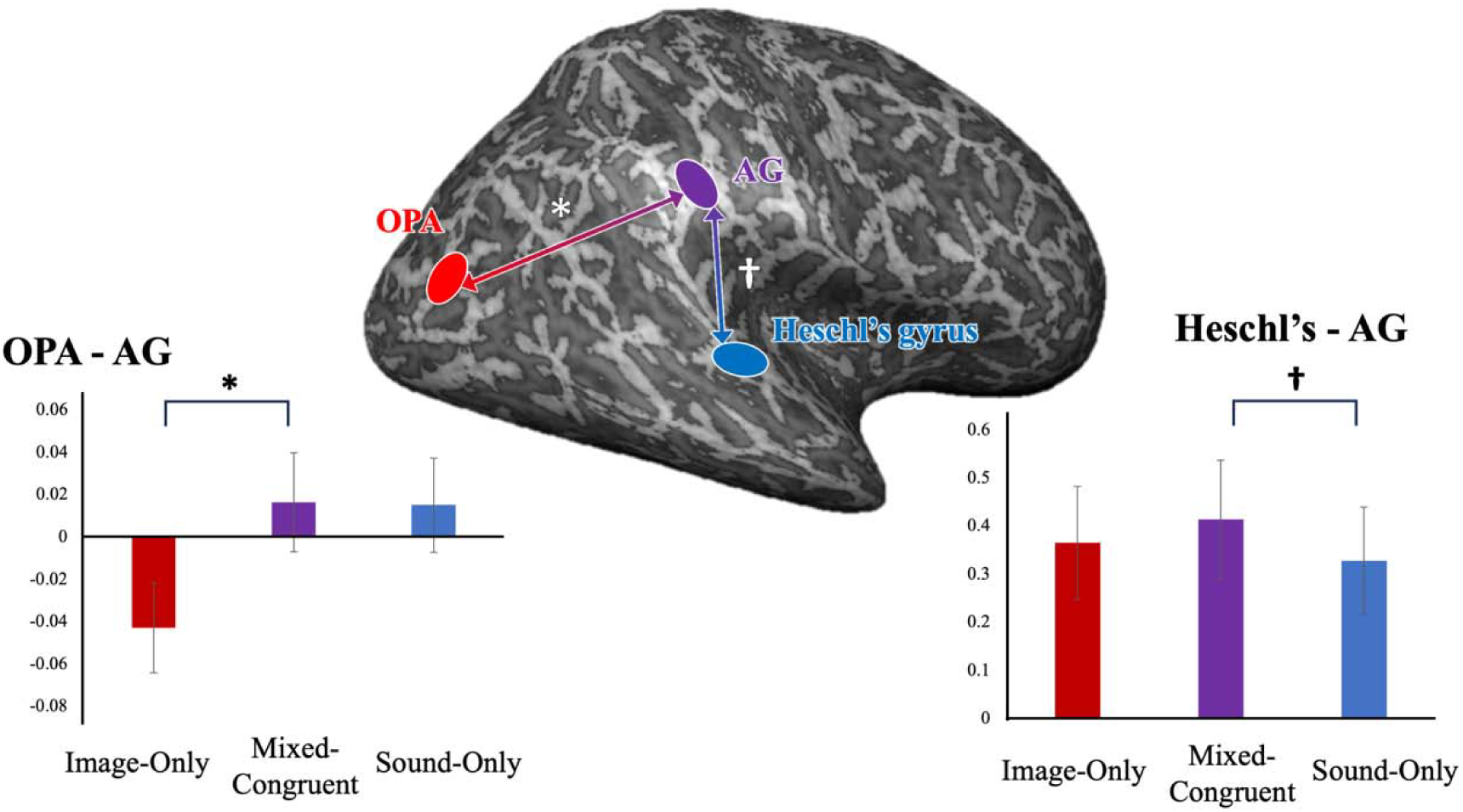
Our analysis mainly examines whether the connectivity between multimodal ROI and modality-specific ROI gets stronger in a multimodal condition over single-modal condition. OPA (marked in red) represents vision specific, Heschl’s gyrus (in blue) audio specific, and AG (in purple) represents modality integrated ROI. Connectivity between ROIs are shown in arrows. Results showed that OPA had statistically significant stronger connectivity with AG when both visual and auditory stimuli were presented (Mixed-Congruent) compared to when only visual stimuli were presented. Correlation between AG and Heschl’s gyrus had noticeable, but not statistically significant connectivity difference when both visual and auditory stimuli were presented compared to when only auditory hstimuli were presented.

## DISCUSSION

This study explored various types of spatial size representation across the human brain. The data confirm previous literature’s findings that patterns of activity in visual scene-specific regions could discern large and small spaces (Cichy & Teng, 2017; Park et al., 2015). We also found supporting evidence that information about the size of space from sounds with different reverberation are reflected in the primary auditory cortex (Norman-Haignere et al., 2013; Teng et al., 2017). The novelty of this study is in testing the two modalities together, asking how the brain represents concurrently presented multi modal information about the size of space. Our results demonstrate that several areas in the temporal, parietal (STG, SPG, STS, AG, MTG, and IPS) and frontal cortex (right IFG, and bilateral SFG) contain audio-visual information about the size of indoor spaces. These areas processed information from eyes and ears separately and together, a representation we call multimodal. Such findings are in line with previous literature suggesting multisensory representation in these areas. For instance, STG, located in the temporal lobe containing the auditory cortex, is known as the site of multisensory integration (Squire, 2009). AG is also known to play a role in in the integration and multimodal perception of sensory inputs (Schendan, 2012; Sheldon et al., 2014), in addition to its role as a cross-modal hub for comprehending and integrating multisensory information (Seghier, 2013).

The term “multimodal” can be interpreted in various ways, depending on the emphasis placed on different characteristics of representation. Information from various sensory organs can simply coexist, like our modality-coexistent model. We can describe this as the ROI reflecting a sum of information, but this does not guarantee their equal weighting of the two modalities. In this study, most of the ROIs showed a preference for visual information over auditory information, a topic we discuss in detail in the following section. Some ROIs are sensitive to the information congruency between modalities, and we define them as modality-integrated representation model. This representation is like the modality coexistent model in that the two information exists in its own form, but only shows significant neural patterns when they are conveying the same information. It is important to note that this modality integrated model should not be confused with our modality-independent (supramodal) model, in which neural specificity is restricted to the information itself (e.g., size of space) rather than the type of sensory input. For example, one group of researchers identified a left-lateralized cortical parcel network that shows similar brain response across reading and listening of sentences using Magnetoencephalography (Arana et al., 2020). Several others discovered areas in the brain responsible for the integration of audiovisual emotional stimuli using fMRI (Klasen et al., 2011; Peelen et al., 2010). Literature on symbolic and amodal concepts were found to be encoded in several parietal and temporal lobes including AG and MTG (Fairhall & Caramazza, 2013; Thakral et al., 2022; Viganò et al., 2021). Furthermore, Jung and colleagues (2018) found that cross-decoding of scene categories across different sensory modalities occur in the prefrontal cortex, suggesting a modality-independent semantic category information of scenes. We also searched for regions that can cross-decode between sensory modalities but could not find evidence for supramodal representation for spatial size.

We believe that the absence of modality-independent representation might be due to the complicated nature of sound perception and its interpretation in relation to the size of space. For instance, we used large reverberating sounds as our stimuli for a large space. However, unlike semantic category of an audio clip that is associated with a specific visual scene (e.g., a beach sound), the echoing sound in our stimuli can have numerous interpretations. An echoing sound may have originated from a large space (e.g., an empty parking lot), or from a small bathroom, where the actual physical space is small, but the reflective materials surrounding the room makes it possible for one to hear a similar, echo-y sound. Therefore, it would be crucial to pair audio information with visual to understand the geometric characters of surrounding space, and this tight pairing could, in turn, hinder the formation of a modality-independent size representation. These complicating factors are expected to be dealt with in future studies, where clutter and wall materials should be included as factors that influence our perception of size of indoor spaces. Another reason why modality-independent size representation could not be found may be due to the parametric nature of the size of space. Because size is a character of space that is relative and parametric, it may be impossible to have a representation pattern that exemplifies space as simply “large” or “small”.

Despite the absence of modality-independent areas, our study found areas that show modality-coexistent and -integrated representation of spatial size in the brain using MVPA. Multimodal areas are regions that show different voxel patterns for large and small indoor spaces when image, sound, or the combination of both were presented. While the MVPA method allows us to examine the neural patterns of voxels, the method has limits in answering questions regarding the nature of the multimodal representation on a neuronal level. For example, it is possible that the neurons within the multimodal voxels are a mixture of neurons that each have image or sound specificity and topographically clustered within the same voxel, or a group of “multimodal” neurons that responds to both images and sounds in single neurons. One method that will allow for further investigation about the nature of neurons is the Repetition Suppression (RS) method (Epstein et al., 2008; Weiner et al., 2010). For example, alternately presenting the two single modal stimuli (i.e., alternating between Image-Only and Sound-Only representation of large spaces) would not elicit RS if a neuron within the ROI each respond to only a single modality. However, if a neuron is “multimodal”, representing both image and sound within a neuron, the same condition will elicit RS. Future investigation will allow us to conclude whether the multimodal representation for the size of space is supported by efficient clustering of sensory-specific neurons, or by neurons that can respond to multi-modal sensory sources.

### Bias to Images in Audio-visual Stimuli

Even though many of our ROIs showed multimodal representation of the size of indoor space, the two modalities were not equally weighted within the multimodal representation. Specifically, the multimodal representation sided with visual information when the visual and auditory size cues clashed (e.g., Mixed Incongruent run types). It is natural for people without severe visual impairment to rely more on their sight than hearing to assess the size of their surroundings (Pike et al., 2012). We tend to depend more on the most reliable source of information (Körding et al., 2007; Noppeney, 2021; Shams & Beierholm, 2010), and this study’s task was to think about the spatial size, thus the more reliable source would have been vision than sound. Lots of acoustical cues influence the reflection of soundwaves other than room size, including materials (DeLong et al., 2007; Hausfeld et al., 1982), shape (Arnott et al., 2013; Hausfeld et al., 1982) and direction (Kolarik et al., 2014) of the enclosed space. These complicating factors make it hard for us to discriminate the size of a room from sound alone. Therefore, reverberation input yields unclear information about the size compared to visual input, which would have led to such sensory bias.

Indeed, the preference for images over sounds has been previously examined in many cognitive research such as motion direction paradigm (Alink et al., 2008; Soto-Faraco et al., 2004), information processing (Colavita, 1974; Spence et al., 2012) and even in design research (Schifferstein et al., 2010). One distinct example of visual dominance is the Colavita effect (Colavita, 1974), where participants presented with audiovisual stimuli tended to respond only to visual stimuli. The experiment consisted of three conditions: visual-only, auditory-only and both presented simultaneously, and the task involved responding to each condition (one response for visual, another for auditory, and both for audiovisual condition). While the participants accurately responded in unimodal conditions, they frequently foundered to miss auditory responses in the bimodal condition (Colavita, 1974; Spence et al., 2012).

Another explanation for the sensory bias in our study is related to the nature of the task itself. The visual and the auditory systems each encode their own spatial and timing information, yet they demonstrate distinct advantages and disadvantages that complement each other (Klemen & Chambers, 2012; Michalka et al., 2015). Our visual system accurately encodes spatial information starting from the retina to various cortical areas in the brain utilizing visuospatial maps (Groen et al., 2022; Silver & Kastner, 2009). Conversely, the judgment of visual temporal events can be affected by non-temporal factors such as contrast (Rucci et al., 2018), as well as inputs from other sensory modalities (Shams et al., 2000). In contrast, the auditory system accurately represents timing information of sounds in early auditory pathways (Adams, 2006; Carney, 1999) but it lacks the a cortical map representation for auditory spatial information (Kong et al., 2014). The contrasting dissimilarities between vision and hearing imply that distinct brain regions are activated based on the task demand at hand, irrespective of the modality of the sensory input. In fact, fMRI experiments have demonstrated that the process of directing auditory spatial attention may involve the utilization of visual spatial maps in the parietal lobe to generate spatial attention maps that integrate multiple sensory modalities (Kong et al., 2014). Taking these findings into account, it is plausible that the task of this study, which involved spatial imagery, evoked a favourable response towards visual input despite the stimuli being multisensory. The observed preference for visual information in our study may be attributed to the inherent advantage of the visual system in accurately encoding spatial information and the interaction between different brain regions involved in multisensory processing.

Although most previous research tackle the issue of the behavioral and cognitive side of visual dominance, we believe the brain’s resolution of our Mixed-Incongruent run types was performed in a similar manner. Humans have natural reliance on their vision, whether it may be because of attentional reasons or just because of how our brain was designed to process multiple stimuli. Regardless, this bias towards sights would have resulted in the Image-Only-data-trained decoder, rather than the Sound-Only, to decipher the data from the Mixed-Incongruent data.

### Background Functional Connectivity and Multimodal Representation

We examined the background functional connectivity between ROIs known for processing visual (V1, OPA, RSC, PPA), and auditory (Heschl’s gyrus) stimuli and modality-integrated representation (AG, right IFG opr) to gain a holistic understanding of the multisensory dynamic of scene perception. Interestingly, we observed that the connectivity between AG and OPA was significantly stronger in the multisensory condition compared to the Image-Only condition. Most other regions, including the marginally increased connectivity between AG and Heschl’s gyrus, turned out to be not salient, suggesting that the connectivity between other sensory ROIs and the multisensory area may not be affected by the input modality. However, it is important to note that previous study using the same method has demonstrated that background connectivity in the ventral visual cortex were linked with the task-relevant semantic category (Norman-Haignere et al., 2012). Moreover, top-down visual attention goals have been found to exert influence on the underlying neural activity between the occipital cortex and category-specific regions such as the fusiform face area and PPA (Al-Aidroos et al., 2012). Such observed difference between previous studies and our current study may stem from the focus on stimulus related changes rather than cognitive state changes. Functional connectivity, particularly background functional connectivity, is often investigated within the context of cognitive process such as memory and attention, rather than stimulus-related aspects. In fact, it has been proposed that measurements based on background functional connectivity are more sensitive to variations in cognitive states, while measurements based on activity are more inclined to reflect distinctions in stimulus-related features of the task in fMRI studies (Li et al., 2023). This may account for the limited findings in our study. However, it does not entirely rule out the possibility that the correlation between ROIs may indeed increase when the stimulus input is multisensory. Future research that manipulates not only the stimulus input but also the cognitive tasks involved with the input would provide further insights into the dynamic changes between brain regions during multisensory processing.

### Early Visual Cortex and Auditory Stimuli

We observed that voxels in the ventral V1 could decode reverberation cues of sounds significantly above chance level. Although seemingly counterintuitive, there are several studies suggesting that early visual cortex could read out complex natural auditory information such as traffic noise and bird singing, in addition to visual sensory information (Petro et al., 2017; Vetter et al., 2014). The neural relationship between vision and audition has been explored in various ways before. For example, Ghazanfar and Schroeder (Ghazanfar & Schroeder, 2006) proffered the idea that the neocortex is multisensory and since then, both ERP and fMRI research provided evidence that even the human primary visual cortex is also considered as multisensory (Murray et al., 2016). Another literature tested whether the divergence between ERP of the ‘sum’ case and ‘simultaneous’ case (Molholm et al., 2002). For the ‘sum’ case, the ERP elicited by visual and auditory stimuli were added, while for the ‘simultaneous’ case, audio-visual stimuli were presented concurrently. Results indicated an audio-visual neural response interaction in the right parietal–occipital area, suggesting the earliest cortical visual evoked potential along with a super additive integration effect. A study using fMRI also demonstrated that the category of natural sounds could be decoded by the neural patterns in the primary visual cortex (Vetter et al., 2014).

Previous literature hypothesized that this multisensory characteristic of the primary visual cortex is because auditory stimulation triggers visual responses (Murray et al., 2016) or a predictive model via feedback from higher multisensory regions (Vetter et al., 2014). However, recent research provided evidence that natural sounds can be decoded from congenitally blind participants’ fMRI activation pattern in the early visual cortex (Vetter et al., 2020), implying that sound decoding in the visual cortex is not purely driven by imagery. Further studies should be conducted to provide an explanation of this peculiar V1’s neural pattern driven by auditory stimuli, but (for the moment) our result that sounds depicting spatial size can be decoded in the primary visual cortex is still aligned with previous evidence.

## CONCLUSION

The current study addresses the neural representation of indoor spatial size when they are visually shown, auditorily played, or concurrently presented. Using multivariate pattern analysis, we found visual-specific processing areas, the primary visual area, OPA, and the right PPA, auditory-specific processing area in the primary auditory regions. Although modality-independent areas were absent throughout the brain, there were several ROIs in the temporal, parietal and frontal cortex that exhibited multimodal representation. We further disentangled the multimodal representation by testing whether the representation contains information about the congruency between the two modalities, specifying the regions that simply have multimodal information coexistent, and the regions that integrate the two modalities and sensitive to the bisensory congruency. Modality-integrated representations in the brain, as well as their interaction with other sensory-specific regions may also provide important and modality-rich information about one’s surroundings, supporting our rapid scene recognition in real-world environments.

## DATA AVAILABILITY STATEMENT

Data for this paper including the visual and auditory stimuli and experimental codes will be made available to download in the corresponding author’s website.

## AUTHOR CONTRIBUTION

Jaeeun Lee: Conceptualization, Methodology, Software, Data curation, Visualization, Investigation, Writing. Soojin Park: Conceptualization, Methodology, Investigation, Supervision, Writing

## ACKNOWLEDGEMENTS

This work was supported by National Eye Institute (NEI) grant (R01EY026042), National Research Foundation of Korea (NRF) grant (NRF-2023R1A2C1006673) to SP. The authors declare no competing financial interests.

**Appendix Table 1.**

| ROI | Left |  | Right |  |
| --- | --- | --- | --- | --- |
|  | Mean | S.D. | Mean | S.D. |
| V1 (bilateral) | 6744.00 | 0.00 |  |  |
| PPA | 5856.00 | 0.00 | 4424.00 | 0.00 |
| OPA | 1064.00 | 0.00 | 2008.00 | 0.00 |
| RSC | 8504.00 | 0.00 | 13928.00 | 0.00 |
| Transverse temporal | 1585.56 | 233.27 | 1206.25 | 159.34 |
| MTG | 14624.94 | 1430.81 | 15823.25 | 1476.80 |
| STG | 16180.19 | 1526.53 | 15454.38 | 1155.32 |
| STS | 12070.06 | 1213.41 | 12952.38 | 1307.01 |
| AG | 2313.31 | 409.20 | 2186.19 | 386.38 |
| SPG | 19294.63 | 1831.57 | 18354.13 | 1666.05 |
| IPS | 5918.38 | 661.62 | 6246.25 | 837.80 |
| IFG opr | 5236.03 | 307.95 | 4646.38 | 250.05 |
| IFG orb | 2309.34 | 110.97 | 2415.50 | 124.42 |
| IFG tri | 4109.41 | 298.59 | 4265.41 | 293.05 |
| MFG | 6765.38 | 672.57 | 7217.69 | 622.50 |
| SFG | 29139.69 | 3250.28 | 28107.44 | 2871.68 |

**Appendix Table 2.**

| ROI | Single Condition decoding |  |  |  | Mixed Congruent decoding | Mixed Incongruent decoding |  |
| --- | --- | --- | --- | --- | --- | --- | --- |
|  | Image Decoding | Sound decoding | Image to Sound | Sound to Image | trained with MC | trained with MC (image-based) | trained with MC (sound-based) |
| V1 | 2.89, 0.01, 0.58 | 2.97, 0.01, 0.57 | -0.01, 0.51, 0.43 | -0.00, 0.51, 0.45 | 3.88, 0.00, 0.58 | 6.73, 0.00, 0.63 | -3.92, 1.00, 0.37 |
| Left PPA | 1.16, 0.13, 0.52 | 2.64, 0.01, 0.56 | 0.02, 0.51, 0.45 | 0.01, 0.51, 0.45 | 0.89, 0.21, 0.50 | 2.69, 0.01, 0.54 | -0.26, 1.00, 0.46 |
| Right PPA | 3.51, 0.00, 0.60 | 3.05, 0.01, 0.59 | -0.02, 0.51, 0.43 | 0.00, 0.51, 0.43 | 1.15, 0.16, 0.51 | 2.97, 0.01, 0.56 | -0.87, 1.00, 0.44 |
| Left OPA | 3.51, 0.00, 0.60 | 3.13, 0.01, 0.60 | 0.01, 0.51, 0.47 | -0.01, 0.51, 0.47 | 0.76, 0.24, 0.52 | 1.92, 0.05, 0.54 | -0.29, 1.00, 0.46 |
| Right OPA | 3.14, 0.01, 0.60 | 2.22, 0.02, 0.56 | 0.00, 0.51, 0.45 | -0.00, 0.51, 0.50 | 2.55, 0.02, 0.55 | 3.94, 0.00, 0.56 | -1.34, 1.00, 0.45 |
| Left RSC | 2.55, 0.02, 0.57 | 3.32, 0.00, 0.61 | -0.00, 0.51, 0.49 | 0.01, 0.51, 0.51 | 1.95, 0.05, 0.52 | 2.49, 0.02, 0.51 | 1.07, 0.64, 0.49 |
| Right RSC | 1.91, 0.04, 0.54 | 3.79, 0.00, 0.61 | -0.02, 0.51, 0.48 | 0.01, 0.51, 0.46 | 2.30, 0.03, 0.55 | 6.84, 0.00, 0.62 | -3.78, 1.00, 0.38 |
| Heschl's | 2.37, 0.02, 0.59 | 5.15, 0.00, 0.66 | -0.02, 0.51, 0.45 | 0.01, 0.51, 0.49 | 5.06, 0.00, 0.71 | 0.26, 0.45, 0.48 | 1.25, 0.64, 0.52 |
| MTG | 4.12, 0.00, 0.62 | 3.17, 0.01, 0.62 | -0.01, 0.51, 0.45 | -0.01, 0.51, 0.44 | 2.77, 0.02, 0.56 | 4.54, 0.00, 0.60 | -1.51, 1.00, 0.40 |
| STG | 3.64, 0.00, 0.62 | 4.61, 0.00, 0.65 | -0.01, 0.51, 0.47 | 0.02, 0.51, 0.45 | 5.59, 0.00, 0.64 | 2.55, 0.02, 0.53 | 0.70, 0.94, 0.47 |
| STS | 4.14, 0.00, 0.60 | 2.95, 0.01, 0.60 | 0.01, 0.51, 0.43 | 0.00, 0.51, 0.46 | 2.01, 0.05, 0.55 | 4.23, 0.00, 0.58 | -1.22, 1.00, 0.42 |
| AG | 3.57, 0.00, 0.58 | 6.42, 0.00, 0.66 | 0.01, 0.51, 0.49 | 0.00, 0.51, 0.47 | 3.92, 0.00, 0.62 | -0.02, 0.54, 0.46 | 2.63, 0.17, 0.54 |
| SPG | 3.66, 0.00, 0.61 | 4.27, 0.00, 0.60 | 0.00, 0.51, 0.45 | 0.01, 0.51, 0.43 | 2.07, 0.05, 0.55 | 5.32, 0.00, 0.60 | -2.06, 1.00, 0.40 |
| IPS | 3.59, 0.00, 0.61 | 2.84, 0.01, 0.59 | 0.02, 0.51, 0.42 | -0.03, 0.51, 0.44 | 3.40, 0.01, 0.59 | 4.75, 0.00, 0.61 | -2.29, 1.00, 0.39 |
| Left IFG opr | 3.93, 0.00, 0.60 | 3.24, 0.01, 0.58 | 0.01, 0.51, 0.47 | -0.00, 0.51, 0.49 | 1.89, 0.05, 0.54 | 3.45, 0.00, 0.58 | -1.38, 1.00, 0.42 |
| Right IFG opr | 3.02, 0.01, 0.59 | 2.73, 0.01, 0.57 | 0.01, 0.51, 0.45 | 0.00, 0.51, 0.45 | 4.38, 0.00, 0.57 | 1.31, 0.12, 0.50 | 1.14, 0.64, 0.50 |
| Left IFG orb | 3.30, 0.01, 0.56 | 3.14, 0.01, 0.61 | 0.02, 0.51, 0.44 | 0.02, 0.51, 0.47 | 1.90, 0.05, 0.54 | 1.30, 0.12, 0.50 | 1.52, 0.48, 0.50 |
| Right IFG orb | 2.49, 0.02, 0.56 | 2.57, 0.01, 0.58 | -0.00, 0.51, 0.49 | -0.01, 0.51, 0.53 | 2.57, 0.02, 0.56 | 3.02, 0.01, 0.57 | -1.00, 1.00, 0.43 |
| Left IFG tri | 3.40, 0.00, 0.62 | 2.39, 0.02, 0.58 | -0.01, 0.51, 0.47 | -0.00, 0.51, 0.45 | 1.85, 0.06, 0.54 | 2.08, 0.04, 0.52 | 0.41, 1.00, 0.48 |
| Right IFG tri | 2.84, 0.01, 0.58 | 3.41, 0.00, 0.60 | 0.01, 0.51, 0.48 | 0.01, 0.51, 0.49 | 3.95, 0.00, 0.58 | 1.88, 0.05, 0.53 | 0.21, 1.00, 0.47 |
| Left MFG | 1.76, 0.05, 0.52 | 5.27, 0.00, 0.65 | 0.01, 0.51, 0.48 | -0.01, 0.51, 0.49 | 1.57, 0.09, 0.55 | 3.79, 0.00, 0.57 | -1.24, 1.00, 0.43 |
| Right MFG | 1.24, 0.12, 0.50 | 3.91, 0.00, 0.64 | -0.01, 0.51, 0.45 | 0.01, 0.51, 0.46 | 2.52, 0.02, 0.58 | 2.06, 0.04, 0.54 | -0.18, 1.00, 0.46 |
| Left SFG | 4.81, 0.00, 0.66 | 3.18, 0.01, 0.61 | -0.02, 0.51, 0.49 | -0.01, 0.51, 0.47 | 3.23, 0.01, 0.60 | 2.93, 0.01, 0.56 | -0.29, 1.00, 0.44 |
| Right SFG | 4.33, 0.00, 0.63 | 3.45, 0.00, 0.63 | -0.01, 0.51, 0.42 | 0.00, 0.51, 0.44 | 3.95, 0.00, 0.60 | 4.06, 0.00, 0.56 | -0.58, 1.00, 0.44 |

